# Model-Agnostic Neural Mean Field With The Refractory SoftPlus Transfer Function

**DOI:** 10.1101/2024.02.05.579047

**Authors:** Alex Spaeth, David Haussler, Mircea Teodorescu

## Abstract

Due to the complexity of neuronal networks and the nonlinear dynamics of individual neurons, it is challenging to develop a systems-level model which is accurate enough to be useful yet tractable enough to apply. Mean-field models which extrapolate from single-neuron descriptions to large-scale models can be derived from the neuron’s transfer function, which gives its firing rate as a function of its synaptic input. However, analytically derived transfer functions are applicable only to the neurons and noise models from which they were originally derived. In recent work, approximate transfer functions have been empirically derived by fitting a sigmoidal curve, which imposes a maximum firing rate and applies only in the diffusion limit, restricting applications. In this paper, we propose an approximate transfer function called Refractory SoftPlus, which is simple yet applicable to a broad variety of neuron types. Refractory SoftPlus activation functions allow the derivation of simple empirically approximated mean-field models using simulation results, which enables prediction of the response of a network of randomly connected neurons to a time-varying external stimulus with a high degree of accuracy. These models also support an accurate approximate bifurcation analysis as a function of the level of recurrent input. Finally, the model works without assuming large presynaptic rates or small postsynaptic potential size, allowing mean-field models to be developed even for populations with large interaction terms.

**Author Summary:** As one of the most complex systems known to science, modeling brain behavior and function is both fascinating and extremely difficult. Empirical data is increasingly available from *ex vivo* human brain organoids and surgical samples, as well as *in vivo* animal models, so the problem of modeling the behavior of large-scale neuronal systems is more relevant than ever. The statistical physics concept of a mean-field model offers a tractable approach by modeling the behavior of a single representative neuron and extending this to the population. However, most mean-field models work only in the limit of weak interactions between neurons, where synaptic input behaves more like a diffusion process than the sum of discrete synaptic events. This paper introduces a data-driven mean-field model, estimated by curve-fitting a simple transfer function, which works with larger interaction strengths. The resulting model can predict population firing rates and bifurcations of equilibria, as well as providing a simple dynamical model that can be the basis for further analysis.

## 1 Introduction

The brain is one of the most complex systems known to science, which makes the problem of computationally modeling its behavior and function both fascinating and extremely difficult. The computational substrate of the brain consists of billions of neurons, each with tens of thousands of connections spanning neurons from the local region to distant areas [1]. Computational neuroscientists are thus faced with a problem of scale: while the biophysical behavior of the individual neuron is understood from first principles [2, 3], and large numbers of neurons have been modeled phenomenologically as a dynamical system [4, 5], unifying different scales of description remains a fundamental difficulty and the subject of ongoing research [6, 7]. Today, as neuromorphic systems are emerging for computation and robotic control [8–11] and human brain organoids and surgical samples are becoming crucial systems for studying disease states and neuronal computation [12–16], the question of modeling the behavior of large-scale neuronal systems is becoming all the more crucial.

One of the most popular approaches to unifying model scales is to make the mean-field assumption, namely that all the neurons in the population under consideration, as well as their inputs, are independent and statistically identical [17, 18]. This assumption is applicable to a variety of systems, ranging from randomly connected sparse networks [17] to modular networks of biological interest [19]. When connections are not strongly selective, neurons within a population are essentially equivalent and can be represented by one population firing rate. Even networks with all-to-all connectivity can be treated, provided the neurons are stochastic enough to remain approximately independent [20]. Under the mean-field assumption, the members of a neuronal population act as independent samples of a shared statistical distribution; this reduces many problems to the study of the firing rate of a single representative neuron receiving *N* inputs at an average rate *r* as depicted in figure 1. Similar approaches have been used extensively in statistical physics, for example in the study of networks of rate neurons [21] or coupled oscillators [22].

**Figure 1.**
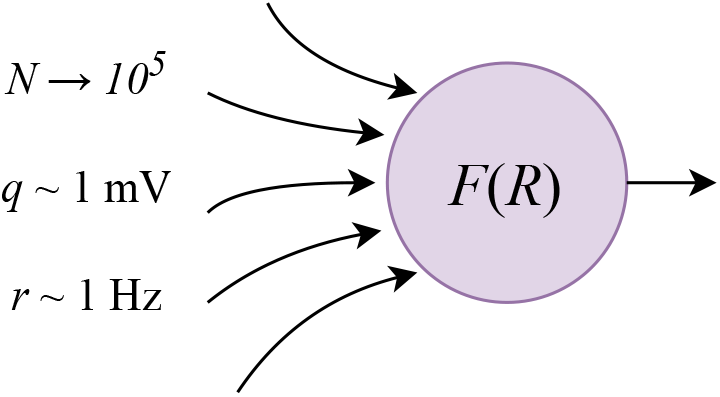
Schematic illustration of a single neuron in a typical network, where the total number of incident presynaptic inputs is large but firing rates and postsynaptic potential sizes are not. The firing rate of the neuron is given by a transfer function *F* of the total input rate *R* = *Nr*.

Many mean-field models examine the limit of an infinite number of presynaptic neurons with infinitesimal postsynaptic potentials, and apply the central limit theorem to replace these inputs with a single white Gaussian noise process. This so-called *diffusion approximation* transforms the single-neuron dynamics into a drift-diffusion problem, for which many techniques have been developed [23, 24]. Perturbation analysis can model complex interactions, but may have difficulty addressing nonlinear phenomena [25, 26]. For more general study of nonlinear effects such as spike frequency adaptation (SFA) or synaptic dynamics, the Fokker-Planck equation can be solved to get the time evolution of the probability distribution of the neuron’s membrane voltage [20, 27]. However, each solution is particular to a single neuronal model, and calculating solutions is computationally expensive even when mathematical analysis has revealed an explicit model [28].

Other mean-field approaches are based on transfer functions, which represent a neuron’s firing rate as a function of some parametrization of the input noise, analogous to the firing rate curve used to study the deterministic dynamics of an individual neuron [18]. Transfer functions do not in principle require the diffusion approximation, but it remains an extremely popular model of synaptic input to transfer functions.

Analytical transfer functions are known for a variety of different neuron types and noise models [29–31], but these require the difficult solution of a first passage time problem and the results are algebraically quite complicated [32].

One line of work in mean-field modeling instead uses a sigmoidal approximate transfer function for a leaky integrate-and-fire neuron in its low-rate Poisson firing regime under the diffusion approximation [17, 33]. This corresponds to the biologically common fluctuating regime of activity, where neuronal firing is asynchronous and irregular [34, 35]. The use of a transfer function additionally requires assuming a stationary distribution of neuronal firing rate [35], but by making a quasistatic assumption, such models can also describe the dynamics of the firing rate [36].

Furthermore, additional variables and parameters can be added to the low-dimensional mean-field system in order to represent other population dynamics such as adaptation variables or population heterogeneity [37, 38].

Mean-field models based on the diffusion approximation and the approximate transfer function of [17] have been applied to a variety of different model neurons [39], have been studied in terms of bifurcation theory [40, 41], and have been compared to biological data [19]. However, the sigmoidal shape of this transfer function does not match analytical results [31, 42]. It also contains an arbitrary timescale *T*, which must be large enough for a quasistatic assumption to hold, but smaller than the minimum interspike interval (ISI) in the system [36]. Furthermore, it is explicitly assumed through moment closure that not only the input but also the membrane potential is normally distributed [43], which is inaccurate for realistic noise sources [44].

In this paper, we propose a novel, explicitly phenomenological approach based on numerically fitting a simple parametrized transfer function that can match both the shape of analytical and observed transfer functions and simulation results. It is also inexpensive to evaluate, as it has only four parameters, which are fitted with a good level of accuracy based on only a few simulations of single uncoupled neurons. Being based on a transfer function, this model is still limited by the assumption of a stationary firing rate distribution, but it makes no assumption about the form or dynamics of the membrane potential distribution. In particular, we do not require the diffusion approximation and can describe populations of neurons with finite numbers of connections and finitely large postsynaptic potentials. We numerically solve a consistency condition to find stable firing rates of a neuronal population, achieving good accuracy in a wide range of simulation examples. We demonstrate the effectiveness of our model by predicting a saddle-node bifurcation to bistability as a function of the connectivity parameter. Furthermore, we implement a popular first-order dynamical model and demonstrate its predictive power alongside its limitations.

## 2 Materials & Methods

There exists a wide variety of dynamical spiking neural models. All are excitable, in the sense that increasing membrane voltage past a threshold causes a rapid return to equilibrium called a spike, but their underlying dynamics vary significantly [45]. We discuss the leaky integrate-and-fire (LIF) neuron below as a representative of this general behavior, but results are given for other models as well.

### 2.1 Leaky Integrate-and-Fire Model

The dynamics of the LIF neuron are given in equation (1). The membrane potential of this neuron undergoes a drift back towards its resting state, while being driven by a time-dependent input *ξ*(*t*). It is also common to introduce a refractory period *T*_ref_, a duration for which the state of the neuron does not change after every spike.

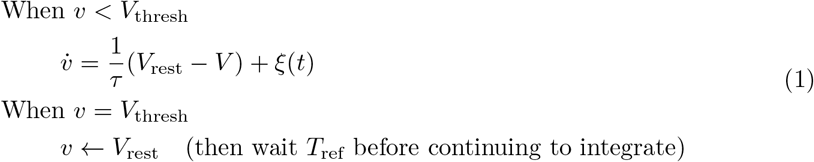

Synaptic input to the neuron is represented by a random process *ξ*(*t*). In this paper, we make the common assumption that this process corresponds to *N* different presynaptic neurons which fire as independent renewal processes at mean rate *r*. Theoretically, such inputs can be treated as a single equivalent Poisson point process with total rate *R* = *Nr* [46]. This assumption is more broadly applicable than it may appear, because according to the Palm-Khinchin theorem, the superposition of *N* independent renewal processes tends in the large-*N* limit to a Poisson process, without requiring that the input renewal processes be Poisson themselves, or even identically distributed [47].

Each presynaptic firing event is a Dirac delta which, when integrated, instantaneously increases the membrane voltage by its fixed postsynaptic potential (PSP) value *q*. A typical order of magnitude for these parameters in the neocortex is 10^4^ neurons [1] firing at a rate of several hertz [17], with PSP magnitudes of about one millivolt [48], as in figure 1.

For neuron models whose implementations do not support delta postsynaptic potentials, alpha postsynaptic currents are used instead. In this case, the weight *w* of a synapse gives the peak value of the postsynaptic current 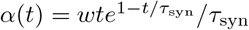, which is integrated by a membrane capacitance *C*_*m*_. To keep inputs comparable between neuron models, we compute *w* as a function of an equivalent PSP size *q* by fixing its integral so that it does not depend on the synaptic time constant and the zero-*τ*_syn_ limit corresponds to a delta PSP of size *q*:

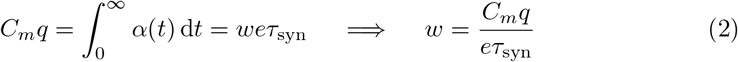

### 2.2 Excitatory-Inhibitory Balance

Although it is convenient to assume that the inputs to the neuron are as homogeneous as possible, models generally assume a distinction between excitatory and inhibitory inputs, which differ at least in the sign of their PSPs. In our simulations, a fraction *η* of the input connections are from excitatory neurons with PSP amplitude *q*_*e*_ *>* 0, and the remainder of the input is inhibitory, with PSP *− q*_*i*_ *<* 0. This applies both to the background input, where the input to each neuron is drawn from two independent Poisson sources with rates *ηR* and (1*− η*)*R* corresponding to its own excitatory and inhibitory background, as well as to simulations with recurrent connectivity, where each neuron is either excitatory or inhibitory and produces the corresponding PSP in all of its postsynaptic neurons.

An exact transfer function is still unknown for even the simple LIF neuron under Poisson inputs [49]. However, it has long been known that in the diffusion limit of infinitely large total input rate *R* = *Nr* and small PSPs, the membrane voltage of the LIF neuron follows an Ornstein-Uhlenbeck process with time constant equal to the membrane time constant *τ* [29]. For a given excitatory fraction *η*, the drift rate *μ* and the diffusion constant *D* of this Ornstein-Uhlenbeck process would be given by:

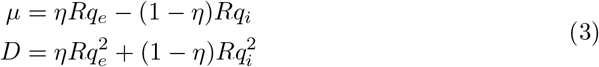

We are interested in modeling neurons in the *asynchronous irregular* state observed in biological systems, which computational experiments have shown occurs when spiking is driven by fluctuations rather than drift [34, 41, 50, 51]. Even where both drift-driven and fluctuation-driven neurons coexist, drift-driven neurons have been described as the predictable background input to the computational activity of fluctuation-driven neurons [52]. The condition where mean synaptic input *μ* = 0, is called *loose* excitatory-inhibitory (EI) balance and is widely observed in experiments and commonly assumed in simulations [53]. Setting *μ* = 0 fixes the inhibitory PSP size *q*_*i*_ as a function of *q*_*e*_:

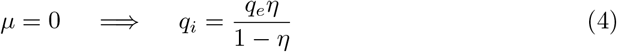

If we then eliminate *q*_*i*_ and define an effective PSP size *q*, the diffusion coefficient can be expressed in terms of it as follows:

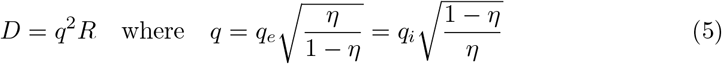

In this paper, we use the value *η* = 0.8, resulting in excitatory and inhibitory PSP sizes that differ by a factor of 4. For example, for *q* = 1 mV, we have *q*_*e*_ = 0.5 mV and *q*_*i*_ = 2 mV. However, the special case where 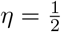 (so *q* = *q*_*e*_ = *q*_*i*_) is also interesting. This corresponds to half of the incoming spikes being inhibitory and half being excitatory, which is convenient mathematically due to symmetry but also has a biological justification. If inhibitory spikes are all mediated by reliable fast-spiking interneurons, their timing would be derived from the same population as that of the excitatory spikes, so the statistics of excitatory and inhibitory spikes would be similar despite the significantly smaller number of inhibitory neurons. It is common to assume something similar in the literature on EI balance in order to achieve mathematically tractable models by avoiding modeling inhibitory neurons explicitly [53, 54].

### 2.3 Refractory SoftPlus

The statistics of interspike intervals (ISIs) in the diffusion limit of equation (3) are found by solving a first passage time problem [29], but these results are mathematically complex, and do not apply to other neuron models, or even outside the diffusion limit (see appendix A).

For this reason, it is appealing to derive a mean-field model from an approximate transfer function that can be fitted numerically to multiple neuron models. In this section, we propose one called “Refractory SoftPlus” after a function occasionally used in machine learning [55]. SoftPlus is a smooth rectified linear function of one variable whose sharpness is controlled by a shape parameter *β*. It generalizes the ramp function *x* ↦ max(*x*, 0), to which it converges in the infinite-*β* limit.

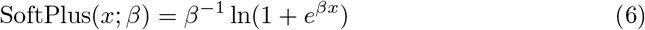

For numerical stability, our implementation behaves as a ramp function for large values of *βx*. The cutoff can be chosen arbitrarily to achieve a desired precision. We use a value of 20, which is accurate to 9 decimal places because ln(1 + exp(20))*/*20 *≈* 1 + 1.03 *×* 10^*−*10^.

To define the Refractory SoftPlus transfer function, we augment SoftPlus <u>w</u>ith scale and shift parameters *α* and *σ*_0_. The input to SoftPlus is taken to be 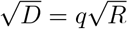 because the firing rate of LIF neuron in the diffusion limit the scales not linearly but with the square root of the diffusion coefficient [29]. We additionally model an absolute refractory period of duration *T*_ref_ by inverting the output of SoftPlus to produce an ISI, adding the refractory period, then inverting again. This yields the following transfer function:

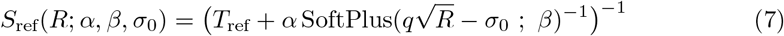

The argument *R* is separated from the three shape parameters by a semicolon to indicate their different roles: *R* is a given input, whereas the shape parameters are derived from fitting to simulation data. Note that *q* appears in the transfer function as a free variable rather than a parameter because it is assumed to be a known constant. For a transfer function which handles variable *q*, see appendix B. For any given neuron model, and for fixed values of *q* and *η*, we compute the firing rate of the neuron over 100 simulated seconds for a range of 500 different values of *R*. For maximum comparability between different values of *q*, we use evenly spaced *R* calculated so that *D* ranges from 0 to 100 mV^2^*/*s. This cutoff point is arbitrary and our results do not depend on it; it is chosen only because it tends to drive our simulated neurons to fire in the tens of hertz, so that most of the shape of their transfer functions can be observed. The four parameters *α, β, σ*_0_, and *T*_ref_ are then optimized to fit this simulated transfer function via nonlinear least squares. To make performance more comparable across models and conditions, we report fitting error normalized by the maximum firing rate observed in the simulation. All simulations were performed using the NEST spiking neural simulator, version 3.3 [56].

Figure 2 shows an example of the performance of this transfer function in comparison to transfer functions from the literature. The simulation data is from the LIF neuron of equation (1), with *V*_thresh_ *− V*_rest_ = 15 mV, *τ* = 10 ms, and *T*_ref_ = 2 ms, subject to Poisson inputs with *η* = 0.8 and *q* = 1 mV (giving *q*_*e*_ = 0.5 mV and *q*_*i*_ = 2 mV). It is compared to the analytical solution for the diffusion limit [29] as well as ReLU and sigmoidal transfer functions. These are simply univariate functions of 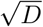 augmented with scale and shift parameters; for the ReLU, this is the ramp function defined above [55], whereas for the sigmoidal transfer function it is the hyperbolic tangent [18]. In each case except the analytical solution, the transfer function is fitted to the left half of the data, i.e. the region where *R <* 50 kHz. The full range of values is then used to show not only the performance of the original fit but also its ability to extrapolate.

**Figure 2.**
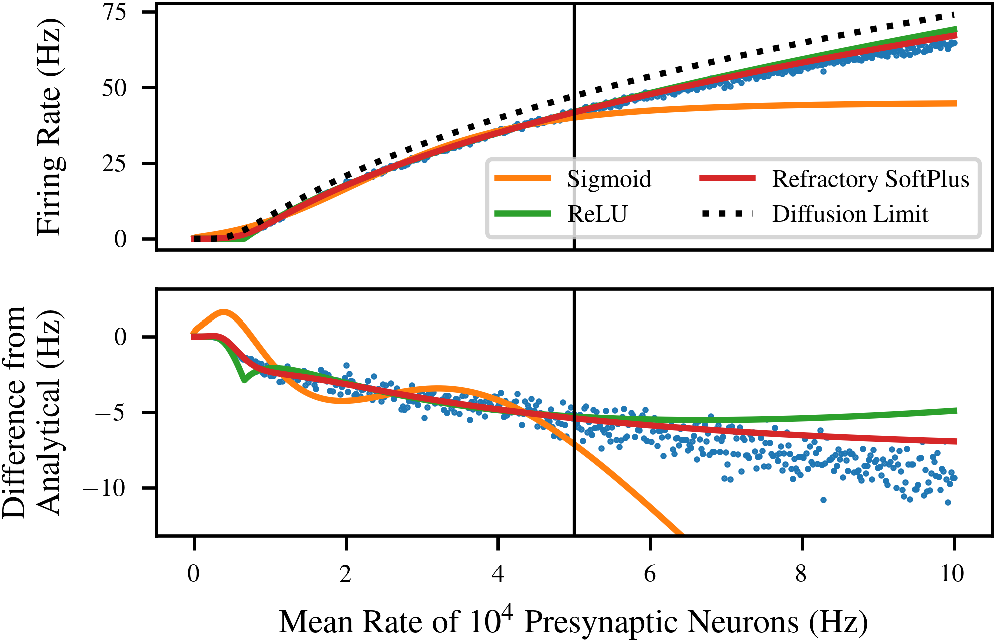
Fit and extrapolation performance of four potential transfer functions for the LIF neuron. Sigmoid, ReLU, and Refractory SoftPlus transfer functions are all fitted to half of the data (left side). Refractory SoftPlus performs well even in extrapolation (right side). Top: the four transfer functions plotted on top of simulated data (blue dots). The sigmoid appears adequate on the training data, but has high error in the extrapolation regime due to saturating. Bottom: the three fitted transfer functions and the simulated data plotted as “error” from the diffusion limit.

The curve for the diffusion limit (black dashed) represents an analytical solution in the limit of infinitesimal PSP size (with constant *D*) and simulation time step.

However, in this simulation, where both quantities are finite, the analytical solution loses significant accuracy, resulting in a significant bias in the reported “error” in the lower panel. The fitted ReLU (green) performs better, but suffers from an overestimate in the extrapolation regime, as well as a small downward bump caused by its sharp elbow. On the other hand, Refractory SoftPlus (red) is capable of capturing the functional form of the simulated transfer function, with virtually no error in the interpolation regime, and lower error in extrapolation as well.

### 2.4 Mean-Field Modeling

Next, we apply the Refractory SoftPlus fit of a neuronal transfer function to derive a mean-field model for the equilibrium firing rate of a recurrently connected population of *M* neurons, each of which fires as a renewal process at the rate *F* (*R*) when subject to a Poisson input with total rate *R*. We assume that each neuron is subject to background synaptic input at a total rate *R*_bg_, in addition to receiving inputs from a random set of *N* other neurons within the population. We exclude multiple and self connections from the present analysis because they could interfere with independence assumptions. The neurons within the population are grouped into an excitatory and an inhibitory subpopulation such that the excitatory fraction *η* is the same for the population as for the background input, and the PSP weights *q*_*e*_ and *q*_*i*_ are set as described in section 2.2 as well.

Under these conditions, the recurrent input and the background input will behave similarly, such that the neuron can be described by its transfer function using a total input rate *r*_eff_ = *R*_bg_ + *Nr*. This enables writing a consistency condition which can be solved to find the equilibrium firing rate of the population:

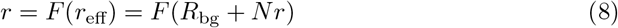

The solutions to this equation are visualized for several concrete values of *N* in figure 3. In this example, we numerically fitted the parameters of equation (7), as described in section 2.3, for LIF neurons with the large PSP size *q* = 5 mV and sparse Poisson background input *R*_bg_ = 0.1 kHz.

**Figure 3.**
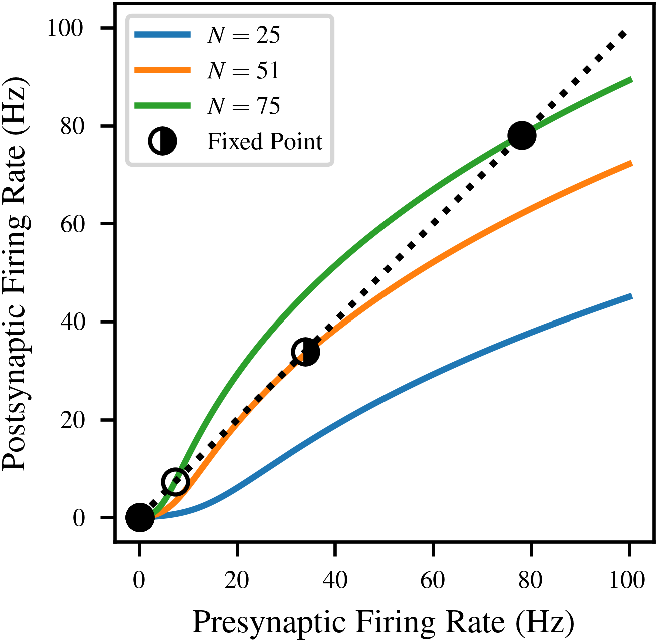
Solutions to the mean-field consistency condition of equation (8) for LIF neurons with *q* = 5 mV and background noise rate *R*_bg_ = 0.1 kHz. Solid curves correspond to different amounts *N* of recurrent connectivity, and give the postsynaptic firing rate as a function of the presynaptic firing rate; the mean-field system is at equilibrium when both are identical (black dashed line). The stability of equilibria is indicated by the fill of the marker (stable = filled, unstable = unfilled, half-stable = half-filled). The model predicts a saddle-node bifurcation, where sufficient levels of recurrent connectivity result in a bistable system.

As shown in the figure, the number and location of solutions to the consistency condition depend on the value of the connectivity parameter *N*. When there is very little recurrent connectivity, the only stable firing rate is the spontaneous rate caused by the background input, very close to *r* = 0 (blue). As *N* increases, the consistency curve gradually slopes upwards to contact the *r* = *F* (*r*_eff_) line. This causes a saddle-node bifurcation at *N* = 51, which creates a half-stable fixed point at *r* = 33.9 Hz (orange). As *N* continues to increase, the bifurcation continues, with the half-stable fixed point becoming one stable and one unstable fixed point (green).

Note also that the background input rate *R*_bg_ can be quite large compared to physiological firing rates (even in the tens of kHz) because it represents the combined activity of a large number of input neurons. In some cases it is also notationally convenient to break *R*_bg_ down into a background input population size *N*_bg_ and mean rate *r*_bg_ such that *R*_bg_ = *N*_bg_*r*_bg_.

### 2.5 Dynamics Model

So far we have described only the equilibrium firing rate of a population of neurons, but it is typically of greater interest to investigate the response of a population to a time-varying external input. We do this using a simplified version of the Master Equation approach [36]. As in the prior work, we assume that the mean firing rate evolves according to linear dynamics with a fixed characteristic timescale *T*, but we only model dynamics to the first order, i.e. we do not model the covariance of multiple interacting populations [35]. This yields the following simple dynamical model:

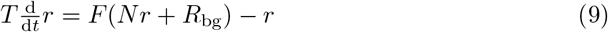

The dynamics of equation (9) have stationary points at the zeros of its right hand side, which are exactly the fixed points of equation (8). In prior work on the master equation formalism, the timescale was considered somewhat arbitrary, but its value affected the parameters and maximum firing rate of the transfer function [36]. Even in previous data-driven mean-field approaches, this necessitated careful selection of the timescale *T*, large enough that the dynamics are memoryless but smaller than the minimum ISI in the system in order to not limit firing rate due to the sigmoidal shape of the transfer function [35]. However, in our model, *T* appears nowhere but in the dynamics, so it must be large enough to enable the quasistatic assumption, but is not limited by firing rate. This in turn allows the model to retain accuracy even in the presence of higher-order dynamical features such as spike frequency adaptation (SFA), so long as the phenomena of interest in modeling have a slower timescale than the phenomena which must be averaged out.

## 3 Results & Discussion

### 3.1 Transfer Function Fit Performance

In order to assess the practicality of a data-driven mean-field model based on a Refractory SoftPlus fit, we first considered the amount of simulation data required to achieve an accurate fit. The computation time required to simulate a neuron’s transfer function scales as *O* (*ST*), i.e. bilinearly with the number *S* of different input rates considered and the total simulated time *T* over which firing rates are calculated.

Increasing either of these quantities increases the quality of the fit by increasing the precision or number of data points used. However, the discrepancy between Refractory SoftPlus and the true transfer function of the neuron also imposes a limit on the achievable accuracy. Therefore, we sought to minimize the computational costs of achieving the best possible quality of fit to the simulated transfer function.

Figure 4 demonstrates convergence of the fit for the LIF (blue), Izhikevich (orange), and the Hodgkin-Huxley (green) neurons. For each model, we simulated the transfer function once for large numbers *S* of input rates and once for large *T*, then considered subsets of the data. Fits were performed to subsets of the full simulation output, and the error of the fitted function was assessed on the final simulated transfer function. The PSP amplitude was fixed at *q* = 1 mV, and input rates ranged from 0 to *R* = 100 kHz.

**Figure 4.**
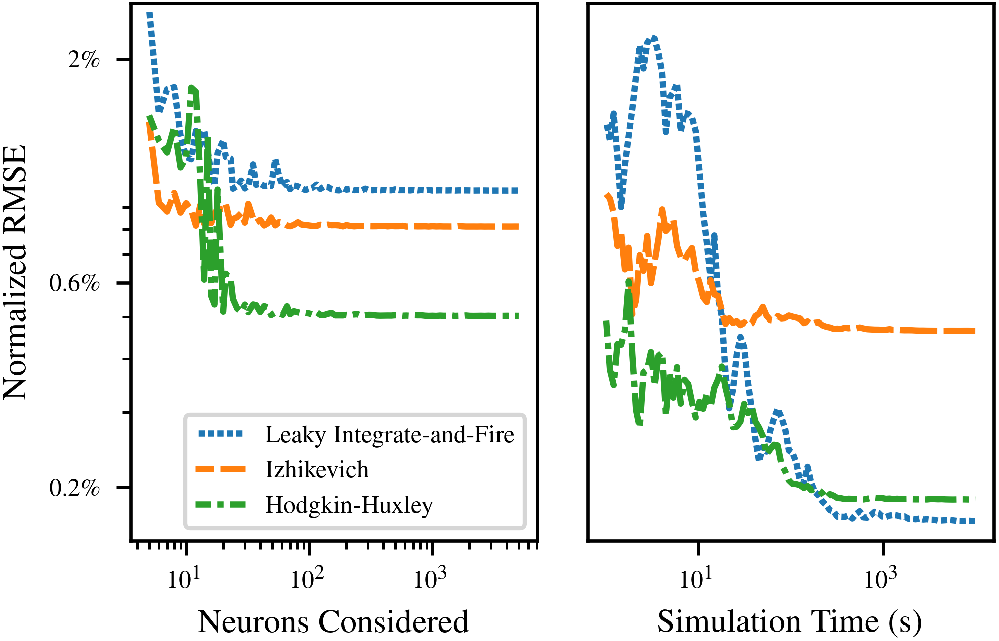
Left: convergence of relative RMS error in the SoftPlus approximation with the number *S* of input rates sampled at a fixed simulation duration *T* = 100 s. The simulated transfer function for the full simulation is compared to Refractory SoftPlus functions fitted to subsets of the data. Right: convergence for *S* = 100 different rates sampled with increasing sample duration *T*.

The left side of figure 4 demonstrates convergence with the number *S* of input rates sampled. The neuron was simulated for a total duration *T* = 100 s at *S* = 10^4^ separate values of the input rate. Fits were then calculated for evenly spaced subsets of this range. The fit error is already only a few percent at *S* = 5, and decreases to a final plateau value by *S* = 100. This can be attributed to the small number of parameters included in the fitted transfer function, which allows it to match observations without a large number of different data points.

The right side of figure 4 shows the convergence of the fit error as the duration of simulation considered for the calculation of firing rate for each neuron increases from *T* = 1 s to *T* = 10^4^ s. A single simulation was performed of *S* = 100 different rates for a duration *T* = 10^4^ s. Shorter simulation times were handled by considering only the beginning of the simulation data. This ensures that any startup transients and nonstationary effects, e.g. due to SFA, are included to the same extent that they would be in disjoint simulations. Again, fit error begins at only a few percent, and inclusion of more data in the fit soon reaches a point of diminishing returns.

Interestingly, for smaller *T*, the error of the fitted curve was substantially lower than the error of the short-time firing rate estimates to which the curve was fitted. The advantage continues for *T* up to 2.7 *×* 10^3^ s for LIF neurons, 1.1 *×* 10^2^ s for Izhikevich, and 7.4 *×* 10^2^ s for Hodgkin-Huxley, with error below 0.5% in each case. (After this point, the error in the short-time firing rate estimates continue downwards as *T* ^*−*1*/*2^ as suggested by appendix C.) This demonstrates that the fitted Refractory SoftPlus transfer function is able to reject a significant amount of noise in the empirical transfer function by providing averaging between adjacent input rates.

### 3.2 Generalization to Other Neuron Models

In the majority of this paper, we use the simulator’s default parameters, but varying these does not affect our conclusions. Indeed, the results can be generalized not only to other parameter values, but also to other neuron models. To demonstrate this, we assessed the quality of the fit as the parameters of each neuron model were varied randomly according to the distributions given in table 1. In each case, dynamics of a single postsynaptic neuron were numerically simulated for 100 seconds subject to balanced Poisson input with *η* = 0.5 and *q* = 1.0 mV at 500 different total input rates *R* from 0 to 100 kHz.

**Table 1:**
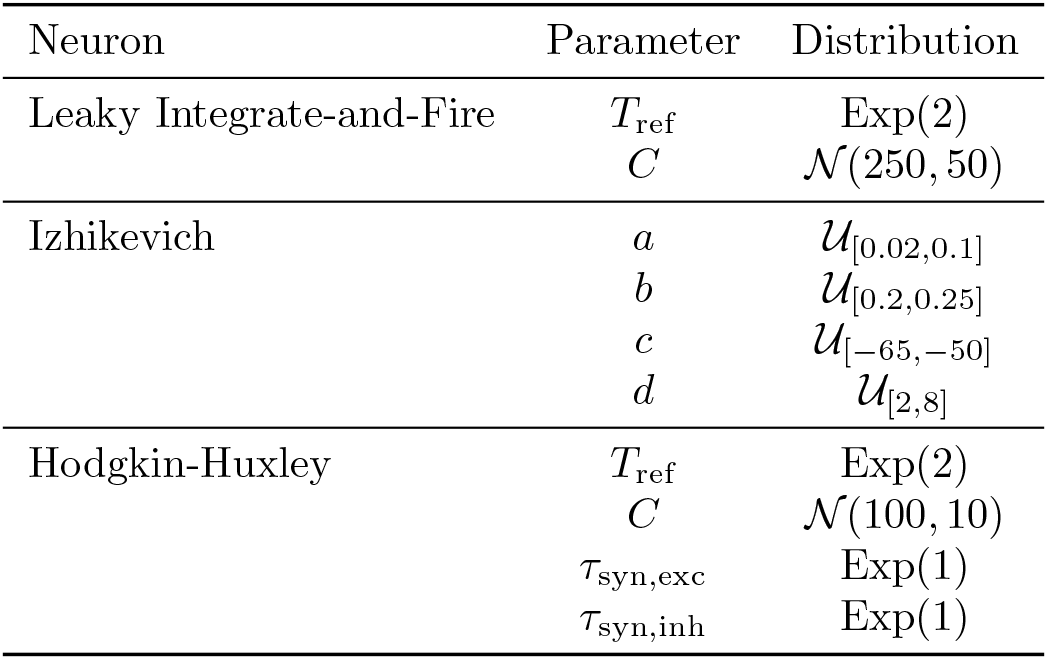
Distributions of random parameter values used to generate figure 5. The notation Exp(*μ*) represents an exponential distribution by its mean, *N* (*μ, σ*) represents a normal distribution by its mean and standard deviation, and *U*_[*a,b*]_ represents a uniform distribution over an interval.

| Neuron | Parameter | Distribution |
| --- | --- | --- |
| Leaky Integrate-and-Fire | $T_{\text{ref}}$ | $\text{Exp}(2)$ |
| | $C$ | $\mathcal{N}(250, 50)$ |
| Izhikevich | $a$ | $\mathcal{U}_{[0.02, 0.1]}$ |
| | $b$ | $\mathcal{U}_{[0.2, 0.25]}$ |
| | $c$ | $\mathcal{U}_{[-65, -50]}$ |
| | $d$ | $\mathcal{U}_{[2, 8]}$ |
| Hodgkin-Huxley | $T_{\text{ref}}$ | $\text{Exp}(2)$ |
| | $C$ | $\mathcal{N}(100, 10)$ |
| | $\tau_{\text{syn,exc}}$ | $\text{Exp}(1)$ |
| | $\tau_{\text{syn,inh}}$ | $\text{Exp}(1)$ |

This avoids the difficulties associated with analytical transfer functions, while still achieving a good fit to simulated data, as shown in figure 5. Performance is evaluated on the LIF neuron discussed above, together with the efficient quadratic integrate-and-fire neuron of Izhikevich [57] and the biophysical Hodgkin-Huxley neuron [2]. Crucially, Refractory SoftPlus is a good fit for the transfer function of all these models, despite their different dynamics and qualitative behavior.

**Figure 5.**
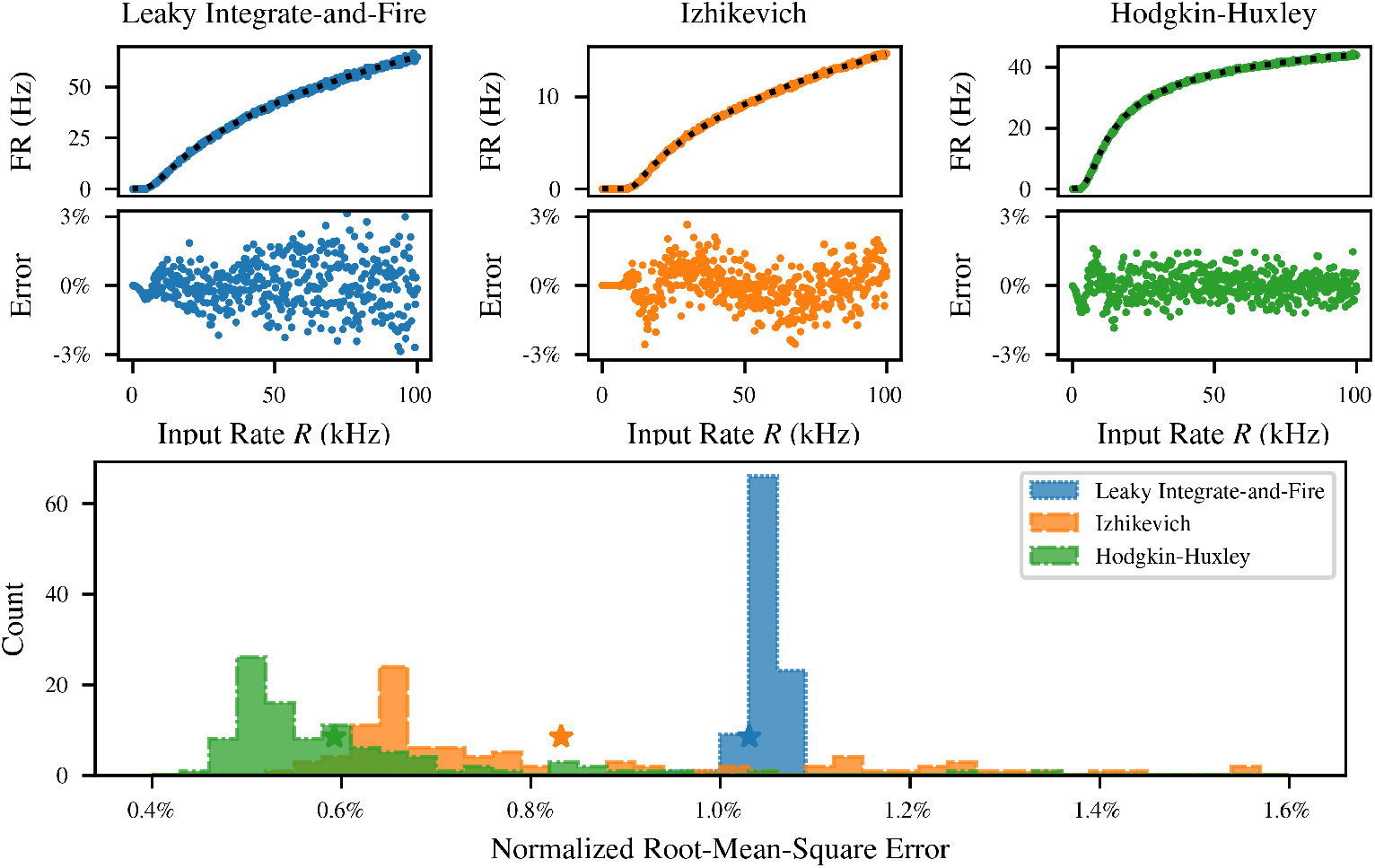
Performance of Refractory SoftPlus fits across neuron models and parameter values. Top: comparison of fits to empirical transfer functions of *M* = 100 LIF (left), Izhikevich (middle), and Hodgkin-Huxley (right) neurons with default parameters, subject to Poisson inputs for *T* = 100 s with *q* = 1 mV. Bottom: distribution of the normalized residual of the fit over 1000 randomly parametrized instances of each neuron. Error for default parameters is indicated with a star.

For the LIF and Hodgkin-Huxley neurons, the distributions were derived from the simulator defaults by giving each time constant an exponential distribution with its default value as the mean, while the membrane capacitance was normally distributed about its default. For the Izhikevich neuron, the distribution of each parameter was made uniform over the range of values considered in the original description of the model [57]. These choices of distribution are arbitrary, but they cover a broad parameter range for each model. For each of the models shown in figure 5, the fitting error under default parameters is indicated with a colored star in the histogram. This point is not an outlier, indicating that our model generalizes well across neuron models for a broad range of parameters.

Also note that for simplicity, the notation with which equation (2) describes the parameterization of synaptic weights to match results between PSP-based and PSC-based neurons ignores the fact that excitatory and inhibitory PSCs may have different parameters. In our simulations, not only were the PSP amplitudes *q*_*e*_ and *q*_*i*_ different because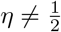but also the synaptic time constants in the Hodgkin-Huxley neuron differ for excitatory and inhibitory synapses. However, despite these asymmetries and significant deviations from the input regime in which the approximations were derived, the flexibility of our assumptions has allowed fits to the simulated transfer function of the Hodgkin-Huxley neuron to be quite good as well.

### 3.3 Bifurcation Analysis

Next we compared the bifurcation and equilibrium locations between the mean-field model and simulation. Activity fixed points were calculated for three different networks of *M* = 10^5^ LIF neurons, as shown in figure 6. Simulations were performed at a range of values of *N*, each corresponding to a different network where each neuron receives connections from *N* presynaptic neurons sampled without replacement from the remainder of the population. All recurrent synapses included an axonal delay chosen uniformly at random between 1 and 15 ms to represent the possibility of neurons spread out in space, but results were similar for small fixed delays as well.

**Figure 6.**
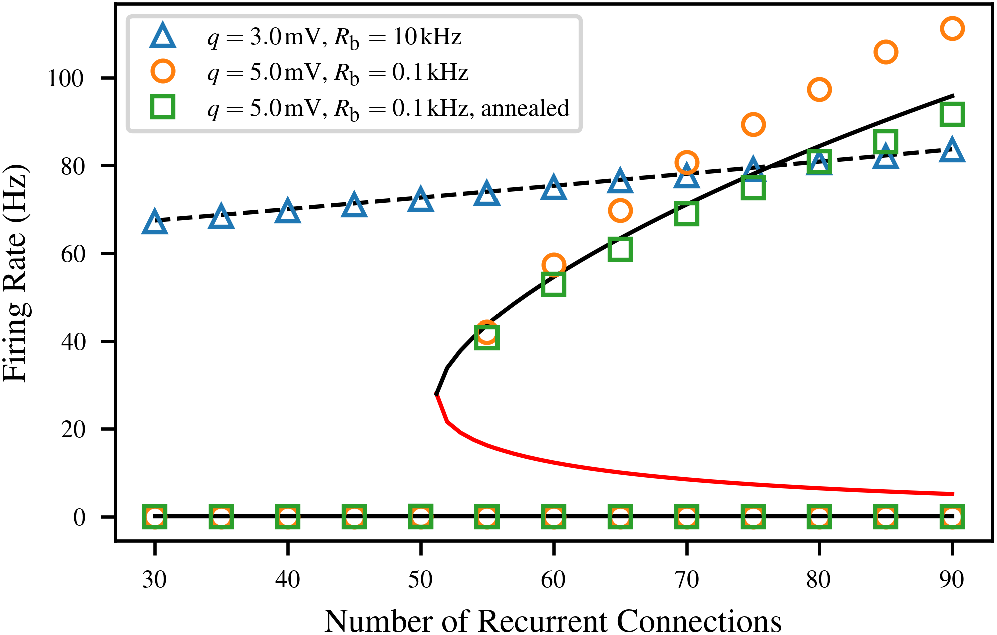
Agreement between numerical solutions to the consistency condition (lines) and observed equilibrium firing rates (markers) for *M* = 10^4^ LIF neurons under three different combinations of connectivity parameters. For the monostable system, the model (black dashed line) and simulation (blue triangles) agree. For the bistable system, the discrepancy between the model (black solid line) and simulation (orange circles) is likely due to temporal correlations between the input neurons, as it can be eliminated by using annealed-average connectivity (green) The unstable fixed points are indicated with the red line.

Each displayed fixed point is the average firing rate across 10 simulations, each 2.5 s in duration. The lower stable population rate is computed by simply simulating the network from a zero initial condition subject to its background Poisson input. To find the higher stable population rate in the bistable network, we transiently raised the firing rate of the population using a “warm-up” input intended to shift it into the basin of attraction of the upper fixed point. We did this by simulating the population without counting spikes for a “warm-up time” of 1 s while the population was subject to additional excitatory Poisson input. To minimize the effects of this drive on the firing rate, we linearly ramped it down from *R*_bg_ to 0 over that time. This was done for all values of *N*, not only those expected to be bistable.

The first condition pictured is a high-noise case which is monostable for the full range of *N*. Here, *q* = 3 mV, and *R*_bg_ = 10 kHz. In this situation, the single firing rate fixed point increases slightly from the baseline spontaneous activity level at *N* = 0, but no interesting dynamics are observed. The model agrees well with simulation for the full simulated range, which is expected because recurrent input contributes under half of the total (ranging from 17% at *N* = 30 to 42% at *N* = 90). This means that there is enough background input to ensure that the assumptions of the model are satisfied even if the recurrent input deviates from Poisson statistics.

The second condition pictured corresponds to the saddle-node bifurcation depicted in figure 3 (orange), with *q* = 5 mV and *R*_bg_ = 0.1 kHz. In this case, the accuracy of the mean-field model suffers somewhat. The network transitions from monostable to bistable between *N* = 50 and *N* = 55 as predicted, but the equilibrium firing rate in the active state is typically noticeably overestimated. This is likely caused by deviations in the input statistics from those used in calibrating the mean-field model. Across the upper branch of this condition, the fraction of input to the neuron which comes from the rest of the network rather than the Poisson background ranges from 97% to 99%. At the same time, firing rates increase, meaning that the absolute refractory period will make up a larger part of the interspike interval, and deviations from Poisson behavior will be more significant.

This is a rather predictable artefact of using such a small value for *N*, when the large-*N* limit is required for the Palm-Khinchin theorem to guarantee a Poisson input as described in section 2.1. Indeed, the overestimation disappears entirely for annealed average connectivity. In this condition, rather than generating a fixed network where each neuron has *N* fixed presynaptic neurons, all-to-all connectivity is combined with probabilistic synapses, whose transmission probability *N/M* is chosen to match the input rate due to *N* presynaptic neurons on average [58]. This breaks up temporal correlations and noticeably improves model predictions of equilibrium firing rates.

An additional caveat must be made specifically for the bistable networks with *N* just above the bifurcation. At *N* = 55, the unstable fixed point, which marks the boundary between the basins of attraction of the two equilibria, is at a relatively high firing rate. As a result, population activity fluctuations are often sufficient to move the network into the basin of attraction of the low-frequency equilibrium. Therefore, for *N* = 55 in particular, we rejected the simulations where this occurred, so the upper fixed point is based on only 1 data point for the second condition and 5 for the third. We did not observe the reverse basin hopping effect, where the network spontaneously jumps to the higher fixed point, because fluctuations near zero firing rate are much smaller.

### 3.4 Finite Size Effects

Although our mean-field model is based on finite connectivity, it does not explicitly depend on the total number of neurons *M* in the population. However, a sufficiently large *M* is implicitly assumed by requiring the neurons to be independent of each other. In this section, we investigate the effect of varying *M* on the dynamical and steady-state behavior of the model.

The implicit assumption of large *M* is similar to a thermodynamic limit: *M* must be large enough that each neuron receives effectively independent inputs, and therefore is itself independent of other neurons. The required value of *M* also depends on the amount of background input; if *R*_bg_ *≫ Nr*, most input to each neuron is from the background, which is independent by construction, so the value of *M* is not very important.

Indeed, we find that the overall population firing rate has a fluctuating component which varies with *M*. Figure 7 explores this by simulating a monostable network with *q* = 3 mV, *R*_bg_ = 10 kHz, and *N* = 75 for a range of values of *M*. The equilibrium population firing rate does not depend significantly on *M* (although estimates become more consistent with larger *M*). On the other hand, there exist fluctuations in the population rate around that fixed point, which decrease in amplitude approximately as *M* ^*−*1*/*2^.

**Figure 7.**
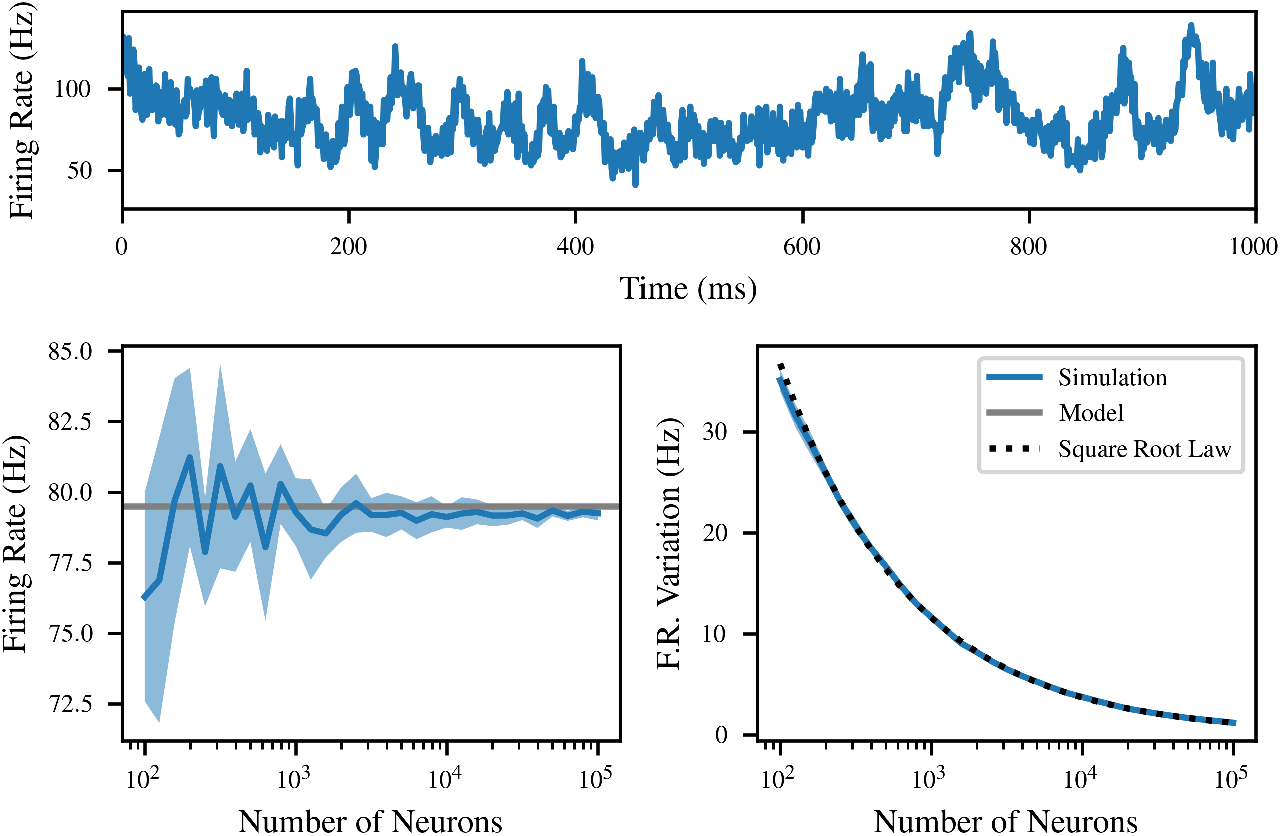
The equilibrium firing rate does not depend on *M*, but time-correlated fluctuations scale as *M* ^*−*1*/*2^. Top: a single example of time-correlated fluctuations in population firing rate at *M* = 10^3^. Left: mean firing rate as a function of *M*, compared to the model prediction (grey). Averaged over 10 simulations, with one standard deviation shaded. Right: fluctuations in firing rate as a function of *M*, exhibiting clear scaling as large populations cancel finite size effects. A fitted inverse square root law is pictured for comparison.

In the figure, ten separate networks with the same parameters were simulated for each value of *M*. The binned firing rate was calculated as the number of spikes in each 1 ms bin of the 2 s recording, divided by *M* as well as the bin size. The result is a time-varying approximation of the firing rate in hertz, which follows a random walk with regression to a mean, as pictured. We quantify this by fitting the parameters of an Ornstein-Uhlenbeck process to the binned firing rate data; the mean is plotted on the left, and the variability 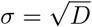 on the right. The average of the mean of this process across multiple simulations is extremely close to model predictions, especially as *M* increases. The variability of the population firing rate within each simulation decreases proportionally to *M* ^*−*1*/*2^, as might be expected from the central limit theorem. This is illustrated using the function *M → kM* ^*−*1*/*2^ with *k* fitted to the data by least squares (black dashed). The variation in the mean between simulations also decreases, but does not follow such a clear functional form.

Particularly surprising is the accuracy of the fixed point even when *M* = 100. In these simulations, almost half of the input is recurrent, and for small *M*, input is highly correlated between neurons because the 75 synapses are chosen from only 100 neurons. However, the location of the firing rate fixed point remains quite close to the model predictions. Evidently, the simple presence of background input on the same order of magnitude as the recurrent input is enough to rescue the assumptions of the model.

However, despite the insensitivity of the equilibrium location to *M*, an important finite size effect appears in the bifurcation analysis of figure 6. The simulations with *N* = 55 are close to a bifurcation, which means that the unstable fixed point marking the watershed between the lower and upper fixed points is 16 Hz. As a result, at *M* = 10^5^, the fluctuations in population rate are large enough that most simulations “fall off” the fixed point within a few seconds of simulation. As a result, a stable equilibrium may not appear stable in practice due to fluctuations.

We define practical stability as the probability that a network will stay near the upper fixed point for the entire duration of the 2.5 s simulation used to calculate the firing rate in the previous section. If practical stability is low for a particular combination of parameters, the state can be considered metastable: the fixed point still exists theoretically, but it may be difficult to observe. This finite-size effect was recently investigated in detail in networks of quadratic integrate-and-fire neurons [59].

Table 2 presents the practical stability of this particular case across a range of values of *M*. There is a clear trend towards higher practical stability as population rate fluctuations are reduced, with better practical stability for annealed-average than for static random connectivity. 50 simulations were performed for each case in the table.

**Table 2:** Practical stability of the upper equilibrium at *q* = 5 mV, *R*_bg_ = 0.1 kHz, and *N* = 55 in figure 6, expressed as the percentage of 2.5 s simulations which avoid dropping to the other equilibrium as a function of *M*. Results are given for static random networks as well as annealed average connectivity.

| Network Size | Stability |  |
| --- | --- | --- |
|  | Static | Annealed |
| $M = 5000$ | 0% | 0% |
| $M = 10000$ | 2% | 10% |
| $M = 20000$ | 10% | 72% |
| $M = 50000$ | 80% | 100% |
| $M = 100000$ | 100% | 100% |

### 3.5 Dynamical Response

Finally, we investigated the ability of the simple dynamics described in section 2.5 to model the sinusoid-following behavior of a recurrently connected network. Figure 8 gives several examples. In each case, a network of *M* = 10^3^ neurons with *q* = 3 mV, *N* = 50, and *R*_bg_ = 10 kHz was simulated subject to independent per-neuron Poisson background input as above, but in this case it is not stationary. Instead, the rate of the background input varies sinusoidally with time at a frequency *f*_bg_ = 1 Hz. Two different amplitudes *A*_bg_ of this oscillation (colmuns) are considered across three different neuron models (rows). For *A*_bg_ = 5 kHz, the background rate fluctuates between 5 and 15 kHz, while for *A*_bg_ = 10 kHz, it fluctuates between 0 and 20 kHz.

**Figure 8.**
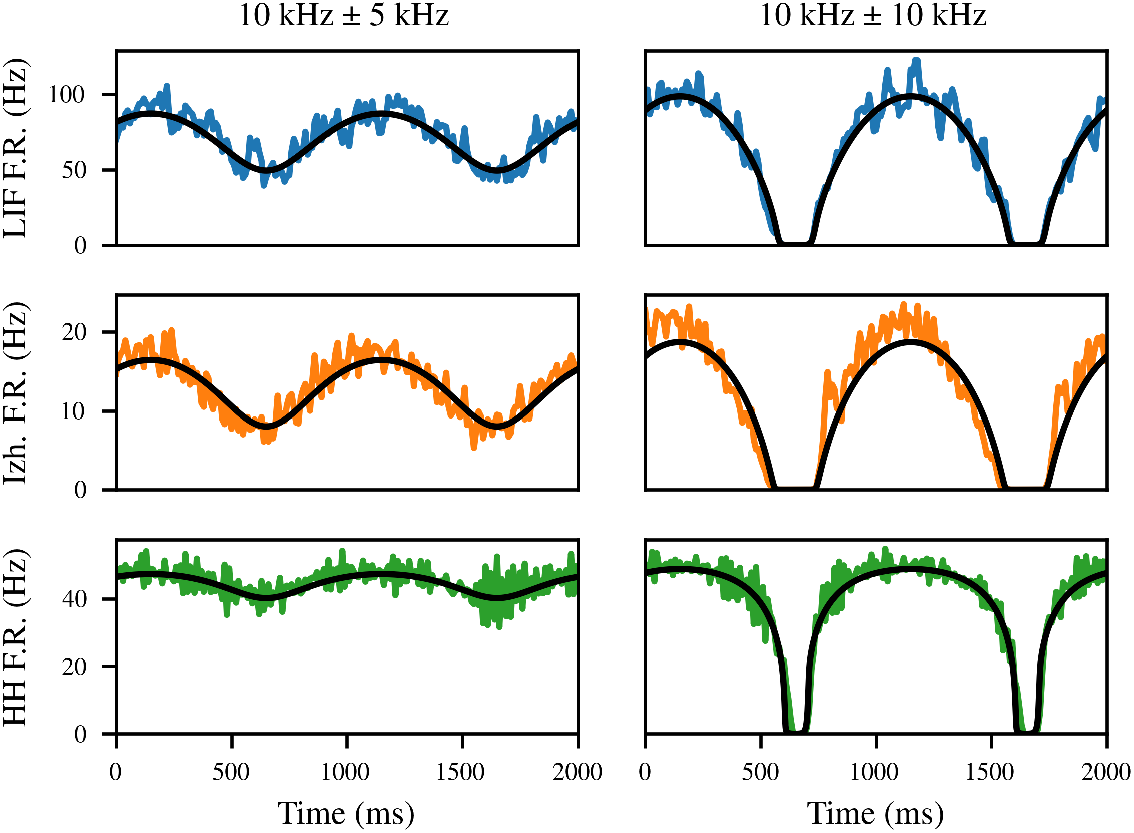
The mean-field model (black) is capable of predicting the dynamical response of recurrently connected networks to a time-varying stimulus. The leaky integrate-and-fire (blue) and Hodgkin-Huxley (green) networks are. modeled quite accurately, whereas the Izhikevich network (orange) changes its firing rate in response to stimulus slightly ahead of the model prediction

This results in quite different qualitative behavior, as in the first case, the population firing rate also oscillates roughly sinusoidally, whereas when the background rate drops to zero, the effects appear much less linear.

In general, the response of the population firing rate to fluctuations in the input is much faster than the response of any individual neuron to those changes [32]. This fact is essential to the theory of tight EI balance, where it explains the use of population rate codes despite their energy inefficiency compared to spike timing codes [53, 54]. We find that although the autonomous population rate fluctuations in figure 7 have a timescale in the tens of milliseconds, the population rate responds to a step input virtually instantaneously (not pictured). Indeed, past work suggests that the perturbation response of the population rate is limited by synaptic, not membrane, time constants [32]. Therefore, although our model does not mathematically require the use of a small value of *T*, we set *T* = 1 ms in these simulations.

Interestingly, the firing rate of the Izhikevich neurons appears to lead the prediction of the mean-field model, as if a derivative term is missing from the true dynamics. The derivative effect disappears under a set of parameters where SFA is absent (not pictured). This suggests that the discrepancy is caused by a failure of the model’s stationarity assumption on short timescales. Indeed, a recent detailed mean-field model of all-to-all coupled Izhikevich neurons observed a similar effect which disappeared when SFA terms were replaced with their population mean [60]. This limitation is not absolute, however. First, the small-deviation example is less problematic, because the population continues firing throughout the simulation in this case, so the adaptation level varies less. Also, the Hodgkin-Huxley neuron has larger-timescale effects, including SFA, and yet is modeled quite accurately in both examples.

## 4 Conclusion

We have developed an approximate mean-field model capable of finding the activity fixed points of a randomly connected network of neurons under the influence of synaptic inputs from other neurons in the network together with background Poisson input. This method is applicable to a wide variety of different neuronal models, and parameter values for the transfer function used in the mean-field approximation can be derived from relatively small-scale and short-duration simulations. The resulting model predicts equilibrium and dynamical population firing rates in a randomly connected network as well as the location of a bifurcation from monostability to bistability.

Unlike previous mean-field models, we do this without making a moment closure assumption. Mean-field models often assume a particular distribution of the postsynaptic membrane potential, generally that it is normally distributed [17, 36, 61]. However, realistic membrane potential distributions can be far from normal [44]. Our approach avoids this by instead assuming that the *input* to the neuron is sufficiently similar to the input used in calculating the original transfer function.

We observed an improvement in model accuracy in one parameter regime when the recurrent input within the population was made more Poisson-like by using annealed-average connectivity instead of a static random network. However, annealed-average connectivity is a strong assumption not typical in modeling; if accuracy in this regime is essential, it would be more useful to fit the transfer function under inputs more similar to those which will later be observed. Although the present work assumes Poisson inputs with loose excitatory-inhibitory balance, this assumption is not fundamental, and simulated transfer functions can in principle be calculated for other input models. This possibility is a key advantage of our approach.

The main limitations of our model are in the simplicity of the dynamics. This extremely simplified master equation does not explicitly model the level of spontaneous activity and so does not provide information about fluctuations [35].

Furthermore, under strong spike frequency adaptation, a deviation from the mean-field predictions can be observed due to a violation of the transfer function’s implicit quasistatic assumption [36]. This has been worked around in the past by adding a dynamical variable to specifically model adaptation [62], an approach which would be equally valid here. Incorporating these two effects into the dynamics is a key direction of future work.

As in results of previous authors, this approximation can be viewed as a bridge between computational neuroscience and machine learning perspectives on neural networks. Given that a similar functional form appears to be conserved across multiple spiking neural models, it is conceivable that it serves a function in biological neural networks as well as in artificial neural networks. In any case, theoretical developments may benefit from the existence of such a simple functional form which is robust across such a wide variety of neuronal dynamics. For our own future work, this simplicity lends itself naturally to adding dynamical terms representing other effects not modeled here, in particular synaptic scaling and plasticity, as well as multi-population interactions.

## Declarations

The authors declare no competing interest. Partial financial support was received from Schmidt Family Futures, the National Human Genome Research Institute, and the National Science Foundation. The funders had no role in study design, data collection and analysis, decision to publish, or preparation of the manuscript.

## Data Availability Statement

This paper contains no primary data. All code is available at

https://github.com/atspaeth/RefractorySoftPlus.

## Supplementary Information

### A. Analytical Transfer Function

The mean inter-spike interval (ISI) for an LIF neuron under the diffusion approximation has been known for a long time [29]. It has a complicated functional form, even after significant simplification using the assumptions that the resting and reset potential of the neuron are identical and that the input has zero mean. Here erfi is the imaginary error function, and the notation !! refers to the double factorial:

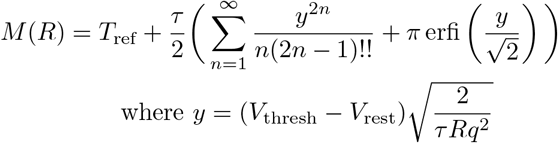

For the LIF neuron, the transfer function is the reciprocal *M* (*R*)^*−*1^. If *T*_ref_ = 0, the transfer function is a smooth curve which is near zero at low *R* but asymptotically grows as *R*^1*/*2^, as shown in figure S1. Also shown in the figure is a neuron with a nonzero absolute refractory period, resulting in a transfer function which grows more slowly with *R*, towards an asymptotic maximum firing rate 1*/T*_ref_.

The neuron was numerically simulated at 500 different input rates *r*. When *q* varies for a fixed *r*, the total number *N* of presynaptic neurons (yielding total input rate *R* = *Nr*) is changed to hold *D* constant so that all conditions are equivalent according to the diffusion limit. Since Poisson processes superimpose linearly, this is only a visual convenience which allows plotting firing rates for different *q* against the same horizontal axis. Firing rates were then calculated by dividing the total number of spiking events by the total simulation time.

**Figure S1:**
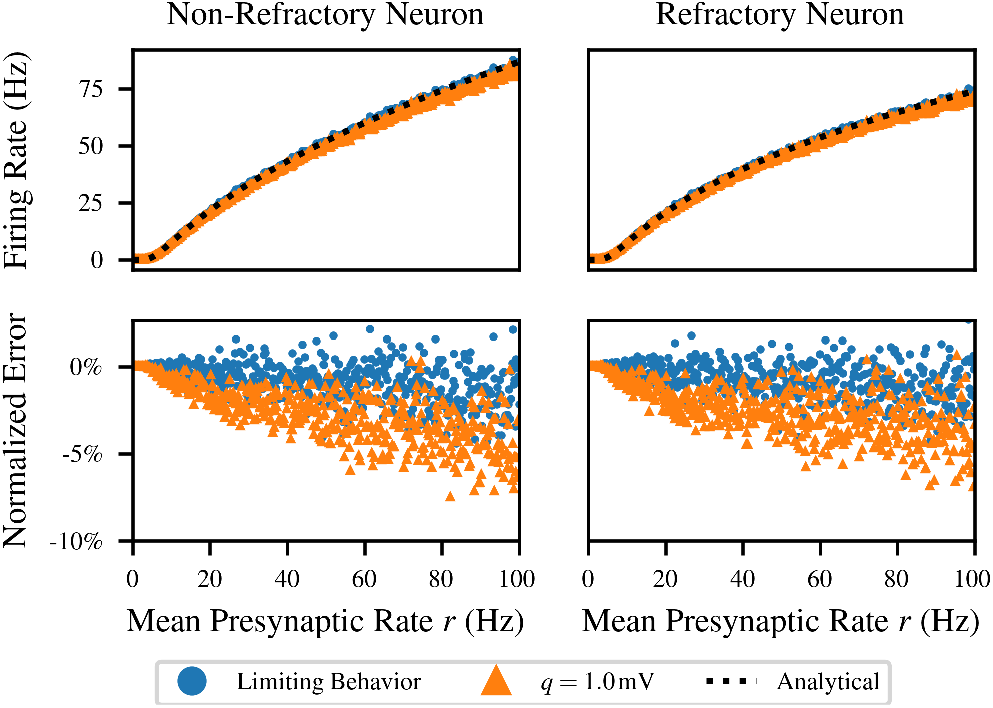
Comparison of the theoretical transfer function *M* (*R*)^*−*1^ to simulated data for an LIF neuron with *τ* = 10 ms, *V*_thresh_ = *−*50 mV, *V*_rest_ = *−*65 mV, and refractory period *T*_ref_ either zero (left) or 2 ms (right). The diffusion limit is represented by a simulation with a small PSP amplitude and large number of presynaptic neurons *q* = 0.1 mV and *N* = 100000 (blue). However, violating the diffusion approximation by changing the PSP to *q* = 1 mV and *N* = 1000 (orange) leads to significant deviation from the analytical solution (dashed line).

**Figure S2:**
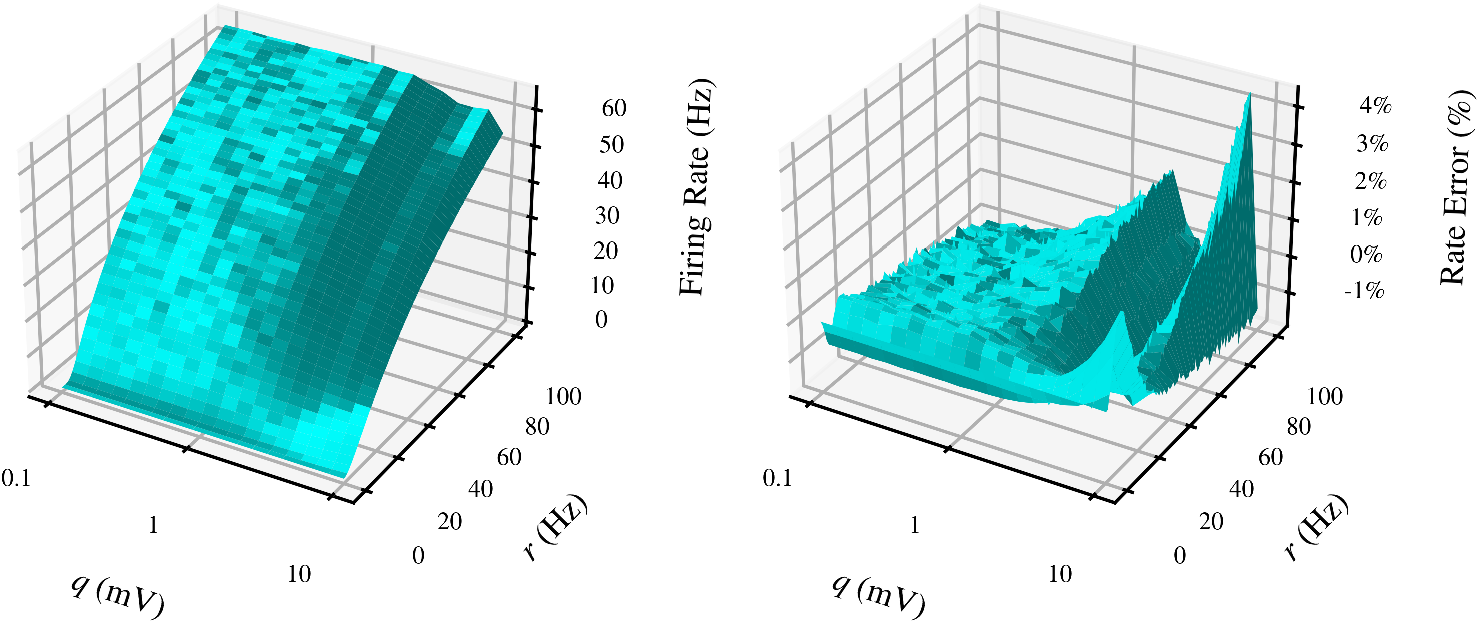
The transfer function as a two-dimensional function of PSP amplitude *q* and presynaptic rate *r* (left), and the error landscape of a single PSP-dependent Refractory SoftPlus fit to this data (right).

Although the analytical transfer function is accurate in the diffusion limit, the simulation in figure S 1 also demonstrates a case where the analytical diffusion approximation breaks down. The limiting behavior is represented by a simulation with a sufficiently small PSP amplitude *q* = 0.1 mV. Increasing the PSP to the biologically reasonable value of *q* = 1 mV (orange) leads to significant deviation from the analytical solution. Similar deviations also occur due to the use of a finite simulation timestep, which leads to a small error even in the better of the two curves in figure S1, as well as to compounded errors from both of these sources in figure 2.

### B. Varying Postsynaptic Potentials

Theoretically, the use of *q* in equation (7) is redundant, as it is equivalent to rescaling *β* and *σ*_0_, but because it is derived from the diffusion coefficient *D*, it acts as a first-order approximation to the effect of PSP amplitude. The only effect of this is to make fitting numerically easier, as the parameters vary less with *q*. However, i t is possible to directly model this dependence to obtain a transfer function with better performance across multiple values of *q*. We observed empirically that *q* has relatively little influence on the fitted values *σ*_0_ and *T*_r ef_, so we mo deled *α* and *β* as having a first-order dependence on *q* with a shared parameter *γ*:

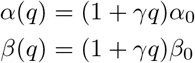

This substitution yields the PSP-dependent Refractory SoftPlus transfer function:

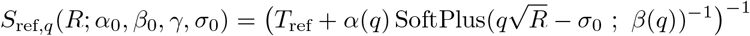

If the simulation data used to fit the transfer function includes varying values of *q*, this version of the function can be fitted instead, which achieves significantly better results using only one additional parameter *γ*. This approach is shown in figure S2, where a simulated transfer function was calculated for LIF neurons subject to Poisson input under a range of *q* and *r*, and a single PSP-dependent Refractory Softplus fit was performed.

**Figure S3:**
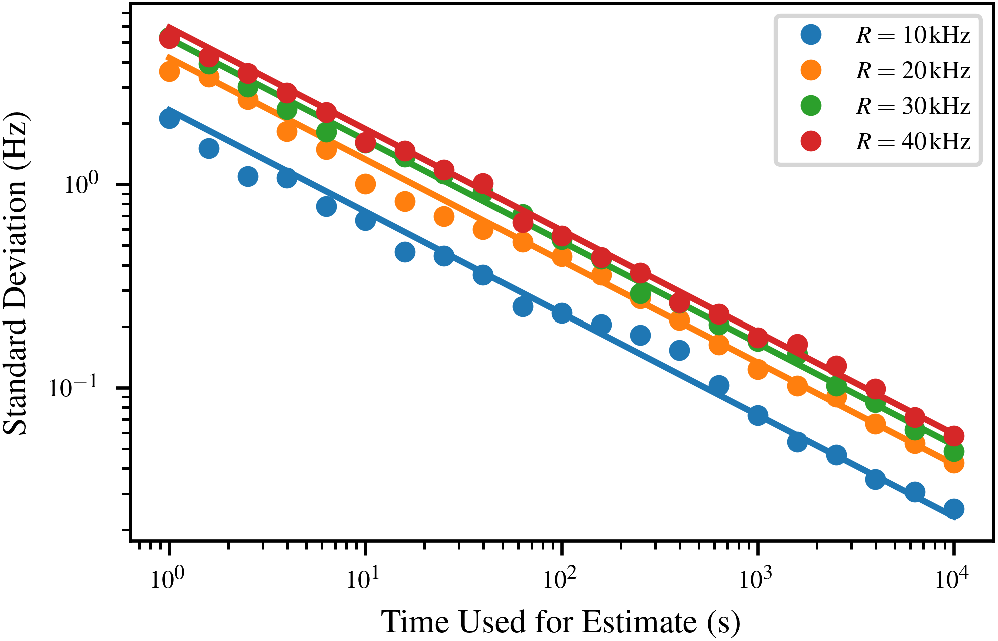
Variance in the firing rate estimate between multiple simulations (colored points) as a function of the simulation time *T*, compared to the approximation (solid lines) for a few different firing rates.

We observe an overall RMS error on par with the simulations of the single neuron, which is able to account for the decrease in firing rate for large *q*. As in figure S1, *N* was covaried with *q* as to keep the range of *D* = *q*^2^*Nr* (equation 3) the same. This choice was made for visualization purposes; fits with a broader range of input *R* and *D* due to unmatched *N* achieved similar performance.

### C. Variance in Firing Rate Estimates

In this section, we provide some context for the magnitude of the error displayed in figure 4 for the convergence of fitted SoftPlus curves as a function of the amount *T* of simulation time available. We do this by comparing the error between the fitted curves and the final long-time simulation run with the error in the firing rate estimates generated from short-time simulations.

If the neuron fires according to a Poisson process with rate *r* over a time interval *T*, the expected number of firings in the time interval is *rT*, with variance *rT*. Therefore, the variance of an estimate of the neuron’s firing rate can be approximated as *r/T*.

We can derive the same result by imagining using bootstrapping to calculate the variance of a binary spike raster. If the neuron fired exactly *rT* times within the time *T*, which is broken up into bins of a very small duration *h*, then each bootstrap fold is *T/h* samples of a Bernoulli random variable with rate *rh*. The sum of these variables follows a binomial distribution with mean *rT* and variance *rT* (1*− rh*) *≈ rT*, so this firing rate estimate also has mean *r* and variance *r/T*. Note that this method assumes that the bins of the raster are independent, which is equivalent to the Poisson assumption.

Figure S3 depicts the standard deviation of 50 firing rate estimates of single LIF neurons with four different input rates. The horizontal axis is the length of the simulation used to calculate the neuron’s firing rate, as in figure 4. Even though the neurons are not actually Poisson, the approximation appears quite accurate for the case considered.

As in figure 4, a full simulation of *T* = 10^4^ s was performed, and data restriction was simulated by restricting attention only to the beginning of this interval. As an artifact of this, adjacent points are correlated in their deviation from the approximation, which we expect would not happen if independent simulations were employed.

In figure 4, the displayed values are the normalized RMS error across an entire empirical firing rate curve. This approximation can be employed to describe the behavior of the error number in the figure, given the true underlying population value of the full sequence {*r*_*i*_} of firing rates of *M* different neurons in the reported population. The expected squared error of each estimate is its variance *r*_*i*_*/T*, so the normalized RMS error *ε* of the entire empirical firing rate curve is given by:

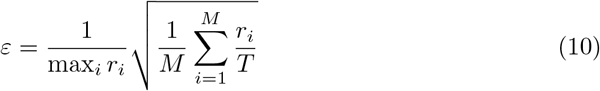

Although the precise value now depends on a significant number of variables which are difficult to know *a priori*, it is at least clear that the expected RMS error will scale as *T* ^*−*1*/*2^ just like the standard deviation of a population of single-neuron estimates.

